# Transposable elements contribute to regulatory hub stress-related long noncoding RNAs in Maize

**DOI:** 10.1101/801704

**Authors:** Yuanda Lv, Fengqin Hu, Yongfeng Zhou, Feilong Wu, Ling Zhou, Brandon S. Gaut

**Affiliations:** Provincial Key Laboratory of Agrobiology, Institute of Crop Germplasm and Biotechnology, Jiangsu Academy of Agricultural Sciences, Nanjing, China; Institute of Soil Science, Chinese Academy of Sciences, Nanjing, China; Department of Civil and Environmental Engineering, UC Irvine, Irvine, CA, USA; Department of Ecology and Evolutionary Biology, UC Irvine, Irvine, CA, USA

**Keywords:** Long non-coding RNA, Transposable elements, Abiotic stress, Co-expression network

## Abstract

Several studies have mined short-read RNA sequencing datasets to identify lncRNAs, and others have focused on the function of individual lncRNA in abiotic stress response. However, our understanding of the complement, function and origin of long-non-coding RNA (lncRNAs) response to abiotic stress, especially transposon derived lncRNA (TE-lncRNA), is still in its infancy. To discover and study lncRNAs in maize (*Zea mays* ssp. mays), we utilized a dataset of 127 RNA sequencing samples that included PacBio fl-cDNA and total RNA-Seq datasets. Overall, we identified 23,309 candidate lncRNAs, 60% of which were identified in polyadenylated (polyA+) samples. The majority (65%) of the 23,309 lncRNAs had sequence similarity to transposable elements (TEs). Most had similarity to long-terminal-repeat retrotransposons from the *Copia* and *Gypsy* superfamilies, representing the high proportion of these elements in the genome, but class II, DNA transposons were enriched for lncRNAs relative to their genomic representation by 2-fold. By assessing the fraction of lncRNAs that respond to abiotic stresses like heat, cold, salt and drought, we identified 1,077 differentially expressed lncRNA transcripts. Their expression was correlated (r^2^=0.48) with their nearest gene, suggesting that lncRNAs are subject to some of the *cis* regulatory features as neighboring genes. By inferring co-expression networks across our large dataset, we found that 39 lncRNAs act as major hubs in co-expression networks, of which 18 appeared to be derived from TEs. These results suggest that lncRNAs, especially TE-lncRNAs, may play key regulatory roles in moderating abiotic responses.

## Background

The functional component of any genome extends beyond its protein coding sequences. Much of the additional function is encoded by RNAs, which vary in size from small RNAs (sRNAs) of< 25 nucleotides (nt) in length, to tRNAs of 70 to ∼90 nt in length, to an even larger class of long non-coding RNAs (lncRNAs). lncRNAs are typically defined as being longer than 200 nt and containing no more than one short (< 100 amino acids) open reading frame [1].

lncRNAs represent a stunning proportion of transcriptional products. In mice, for example, an early study cataloged ∼34,000 lncRNAs, representing one-third of all polyadenylated cDNAs[2]. More recent work has annotated ∼14,000 lncRNAs in humans[3]. Work in plants has lagged somewhat behind, but plant lncRNAs have been identified based on various kinds of high throughput expression data. For example, microarrays have been used to detect 6,480 lncRNAs from *Arabidopsis thaliana*[4];single-stranded RNA sequence data have led to the identification of 2,224 lncRNA transcripts in rice (*Oryza sativa*) [5]; and total RNAseq data have been employed to detect 7,245 lncRNAs in maize (*Zea mays* ssp. *mays*) [6].

At least three general properties of lncRNAs have become apparent from studies of both plants and animals. The first is that many lncRNAs are polyadenylated and capped, suggesting that they are transcribed and processed similarly to mRNAs [7]. However, lncRNAs can also be non-polyadenylated, and hence robust lncRNA discovery requires consideration of both polyadenylated and non-polyadenylated RNA samples. The second is that lncRNAs tend to be expressed at lower levels than coding genes, but with pre*cis*e spatio-temporal patterns [3, 7–13]. A third general property is that some lncRNAs overlap with coding regions and sometimes contain parts of an exon; however, most originate from intergenic spaces (and these are sometimes called long intergenic RNAs or lincRNAs). Consistent with their origin from intergenic spaces, a large proportion of lncRNAs are either derived from transposable elements (TEs) or contain remnants of TEs. For example, Kapusta et al. (2013) determined that 75% of human lncRNAs contained regions that appear to be derived from TEs.

Just as the origin and structures of lncRNAs are diverse, they play similarly varied functional roles. One major role is to act as templates for sRNA production, which in turn often contribute toward the epigenetic silencing of TEs [14, 15]. Some lncRNAs perform other key functions, especially regulatory roles in cellular and developmental processes [3, 16]. In plants, for example, lncRNAs have been shown to affect functions as diverse as phosphate signaling [17], flowering time [18], and susceptibility to pathogens [19]. Consistent with the hypothesis that lncRNAs play important regulatory roles, some lncRNAs are conserved among species and appear to be under purifying selection[3, 20, 21].

A growing body of evidence also points to a potential role for plant lncRNAs in responses to abiotic and biotic stresses. A few studies have identified *Arabidopsis* lncRNAs that respond to salt, drought, heat and cold stresses, as well as phosphate starvation [22–24]. The expression of 28% (1,832 of 6,480) of *Arabidopsis* lncRNAs was found to be significantly altered under biotic and/or abiotic stresses [4]. These findings – i.e., that lncRNAs are associated with stress responses – are particularly important in the context of crop species, because abiotic stresses affect crop yield and quality [13, 25–29]. However, the identification of lncRNAs during crop stress response remains largely unexplored, with a few notable exceptions. For example, 637 nitrogen-responsive lncRNAs and 664 drought-responsive lncRNAs have been identified in maize seedlings [6, 30]. Similarly, 1,010 and 1,503 lncRNAs are known to be differentially expressed under abiotic stress in rice and in chickpea [31].An important but challenging issue is to discover lncRNAs that are associated with abiotic stress responses and then to determine which lncRNAs function as key regulators, which serve as sRNA templates and which represent transcriptional noise.

Here we identify lncRNAs that relate to abiotic stress responses in maize. Our work extends previous maize lncRNA studies in at least three ways [6, 8, 30]. First, our efforts to detect lncRNAs are based on more expansive data. To perform lncRNA discovery, we have amassed 127 RNAseq datasets that were generated by different methods, in different tissues and across developmental stages, with a large subset generated in abiotic stress experiments, including salt, drought, heat, cold, UV and ozone stresses. The data include 89 RNAseq samples based on Illumina sequencing, 36 RNAseq datasets based on PacbioIsoSeq experiments, and two Illumina RNAseq datasets that were based on total RNA to potentially detect non-polyadenylated RNAs. Second, we investigate the relationship between TEs and lncRNAs. More than 85% of the maize genome consists of DNA derived from TEs [32], and we therefore expect that many lncRNAs exhibit similarity to TEs. Thus far, however, the connection between lncRNA and specific TE superfamilies has not yet been investigated for maize. Third, we identify the subset of lncRNAs that are differentially expressed under abiotic stress to begin to narrow the set of candidates that function in stress response. We futher investigate co-expression of lncRNAs with neighboring genes within expression networks to further narrow a candidate list of potentially functional lncRNAs [33, 34]. Bringing these diverse analyses together, we identify several lncRNAs that are hubs in coexpression networks relative to abiotic stress responses, including lncRNAs derived from TEs.

## Results

### Construction of transcripts and lncRNA discovery

To discover lncRNAs and examine their expression during abiotic stress, we used 89 RNAseq samples, 2 total RNA-Seq samples and 36 Pacbio Iso-Seq samples. For the Illumina datasets, we extracted and cleaned ∼305 Gb of sequence data; on average 92.1% of Illumina reads per sample aligned successfully to the maize B73 v4 reference sequence[35]. Aligned reads from each Illumina sample were merged. We also collected and cleaned ∼1.98 Gb of IsoSeq sequences, aligned them to the B73 reference, and collapsed them for a total of 17,673 loci with 43,774 transcripts. We then combined the Illumina RNAseq and PacBio IsoSeq data based on alignment of contigs to the reference, ultimately identifying a non-redundant set of 77,172 loci with 95,523 transcripts (Figure S1). Among these, 19,449 transcripts that were found only in the total RNA sample, representing potential polyA-lncRNAs. The set of 95,523 transcripts consisted of both coding transcripts and potential lncRNA transcripts. To identify the latter, we used a pipeline based on a combination of annotation programs and Pfam analyses (see Methods). Of the 95,523 assembled transcripts, CPC2 annotation identified 31,967 non-coding transcripts (CPC2 score⍰<⍰-1), and 41,839 transcripts were deemed to be noncoding based on CNCI analysis. Of these two sets, 26,099 transcripts were longer than 200 bp and were predicted to be non-coding by both CPC2 and CNCI. These were further filtered by: i) comparing them to the Pfam database, retaining only those transcripts without a match (Blast, Evalue>1e-05) and *ii*) FPKM filtration, based on our requirement that FPKM had to exceed 1 in least one sample. The final dataset, which we consider high confidence lncRNAs, consisted of 23,309 transcripts (Table 1; Figure S1), representing 24% of the total (23,309/95,523). The average length of these candidate lncRNAs was 382 bp. None had an ORF > 100 amino acids in length, as per our definition of lncRNAs (see Methods), but most (95.15%) had one ORF. Among the 23,309 lncRNA candidates, 59.3% (or 13,822 transcripts) were identified from polyadenylated (polyA+) RNAseq samples, and the remaining 40.7% (or 9,487 transcripts) were from total RNA samples, representing potential polyA-transcripts (Table 1; hereafter we refer to lncRNAs from total RNAs as polyA-for simplicity). A file containing all the identified lncRNAs sequences, along with their genomic locations, is provided in DataS1.

**Table 1:** A summary of lncRNA discovery.

|  | Total | %TE-lncRNA | lncRNA | intronic | Fall within<br>annotated<br>lncRNAs | Overlap with<br>annotated<br>lncRNAs |
| --- | --- | --- | --- | --- | --- | --- |
| polyA- | 9,487 | 79.26% | 9,153 | 29 | 189 | 116 |
| polyA+ | 13,822 | 56.37% | 11,346 | 156 | 1,201 | 1,119 |
| Total | 23,309 | 65.69% | 20,499 | 185 | 1,390 | 1,235 |

The 23,309 lncRNAs were widely distributed across the 10 maize chromosomes (Figure S2). We also examined their location relative to annotated coding sequences within the maize genome. As expected from our search strategy, most lncRNAs (87.9%; 20,499 of 23,309) were intergenic, based on the output (a U class code) from gff compare. Only 185 lncRNAs were found to be intronic, with 29 and 156 as polyA- and polyA+ (Table 1). The remaining high confidence lncRNAs corresponded to, or overlapped with, previously annotated lncRNAs in the B73 v4 reference (Table 1). Among the 20,499 lincRNAs, 44.7% (or 9,153 of 20,499) were from total RNA datasets (i.e, potentially polyA-), representing a significant enrichment for lncRNAs within the total RNA samples (Pearson χ-squared; p < 0.001).

### Most lncRNAs are derived from transposable elements

Although a large fraction of lncRNA derive from TEs [7, 36], leading to the hypothesis that TEs have shaped functional domains of lncRNAs [37], this previous paper provided few details about the relationship between TEs and lncRNAs, such as the TE superfamilies that have contributed to lncRNAs or the proportion length of individual lncRNAs that can be attributed to TEs.

To identify which lncRNAs may be derived from a TE, we masked regions of our 23,309 high-confidence lncRNAs using a species-specific TE library (see Materials and Methods). Overall, we found that 65.69% lncRNAs (15,312 of 23,309) overlapped with known maize TEs, which is a proportion similar to the previous maize study based on fewer lncRNAs [8]. Most (61%, or 9,341 of 15,312) TE-lncRNAs showed similarity to TEs over ≥90% of their length (Figure 1A). Perhaps unsurprisingly, the proportion of polyA-lncRNAs that were masked by TE sequence was higher than that of polyA+ lncRNAs (79.26% vs. 56.37%), which is a significant difference (*p*< 1e-5) (Figure 1B). Hereafter we refer to lncRNAs with sequence similarity to TEs as “TE-lncRNAs”.

**Figure 1:**
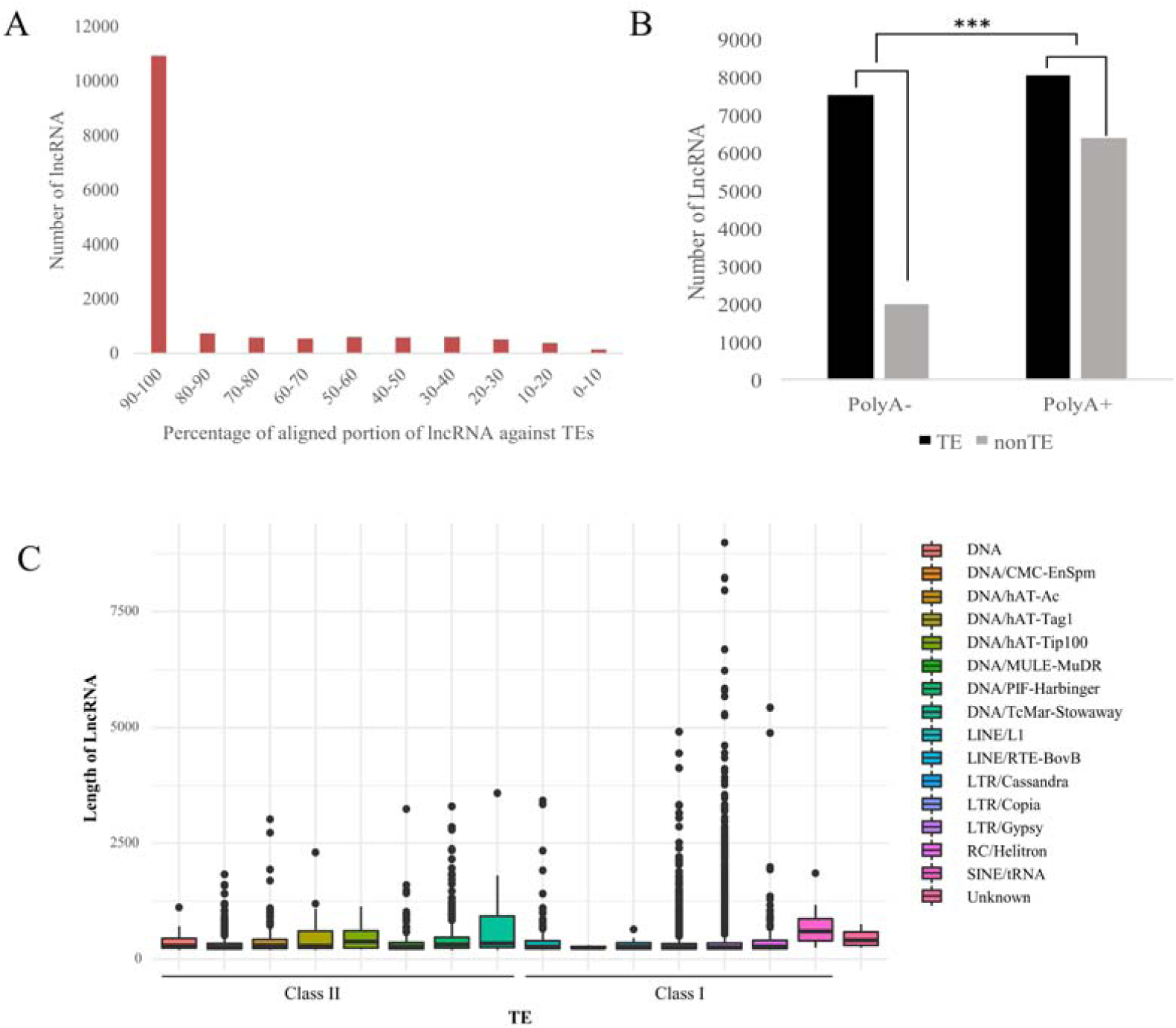
The relationship between lncRNAs and TEs. A) The histogram indicates the number of lncRNAs (y-axis) relative to the percentage length (x-axis) of lncRNAs that have similarity to TEs. B) The numbers of lncRNAs that are polyA-(i.e., from total RNA) or polyA+ with similarity to TEs. The proportion of polyA-lncRNAs is significantly enriched for similarity to TEs. C) The length distribution of TE-lncRNAs organized by their inferred TE superfamily of origin.

We further investigated the superfamily of TEs that were similar to the 15,312 TE-lncRNAs. We found that 86% had sequence similarity to Long Terminal Repeat retrotransposons of the *Gypsy* and *Copia* superfamilies (Table 2) and also that some of these TE-lncRNAs exceeded 3,750 bp in length (Figure 1C). A much smaller proportion of TE-lncRNAs were derived from DNA transposons (Table 2); the longest of these were shorter than the longest TE-lncRNAs with similarity to *Gypsy* and *Copia* elements (Figure 1C).

**Table 2:** The proportion of base pairs attributable to different TE superfamilies based on the total length of inferred TE-lncRNAs and the B73 reference genome.

| TE Class | Number of TE-lncRNAs | % total length of TE-lncRNAs | % total length of TEs<br>in the genome |
| --- | --- | --- | --- |
| **Retroelements** |  |  |  |
| SINE | 15 | 0.36% | 0.02% |
| LINE/L1 | 151 | 1.28% | 0.96% |
| LINE/RTE-BovB | 8 | 0.05% | 0.14% |
| LTR/Cassandra | 47 | 0.23% | 0.10% |
| LTR/Copia | 3917 | 22.45% | 32.39% |
| LTR/Gypsy | 9566 | 61.53% | 59.64% |
| **Total** | **13,704** | **85.90%** | **93.25%** |
| **DNA transposons** |  |  |  |
| DNA | 28 | 0.28% | 0.03% |
| DNA/CMC-EnSpm | 579 | 3.97% | 3.33% |
| DNA/hAT-Ac | 234 | 2.03% | 0.73% |
| DNA/hAT-Tag1 | 20 | 0.56% | 0.08% |
| DNA/hAT-Tip100 | 32 | 0.34% | 0.14% |
| DNA/MULE-MuDR | 213 | 1.70% | 0.93% |
| DNA/PIF-Harbinger | 291 | 2.80% | 0.64% |
| DNA/TcMar-Stowaway | 35 | 0.41% | 0.08% |
| **Total** | **1,432** | **12.10%** | **5.96%** |
| **Helitrons** | 162 | 1.79% | 0.78% |
| **Unclassified:** | 14 | 0.21% | 0.02% |
| **Total** | **15,312** |  |  |

These observations raise an interesting question: Do LTR/Gypsy and LTR/Copia elements give rise to lncRNAs more often than expected, given their proportion of the genome? To address this question, we estimated the proportion length among all annotated TEs that were attributable to LTR/*Gypsy*, LTR/*Copia* and other element superfamilies, based on RepeatMasker analyses. We then compared these percentages to the proportion length among inferred TE-lncRNAs (Table 2). We found, for example, that LTR/*Gypsy* elements produced TE-lncRNAs at roughly the expected proportion (61% vs. 59%), relative to their representation in the genome. However, LTR/*Copia* elements contributed TE-lncRNAs at a lower proportion than their proportion length among annotated TEs (22% vs. 33%). Particularly notable is the fact that class II DNA elements produced TE-lncRNAs in our dataset at ∼2-fold higher rate (12% vs. 6%) than expected based on their total length among TEs in the genome (Table 2).

### Differential expression under abiotic stress

One general feature of lncRNAs is that they are expressed at lower levels than protein coding genes, and they are often expressed tissue specifically [3, 6, 8, 23, 38, 39]. We assessed the expression levels of coding and lncRNA transcripts based on their maximum FKPM across all of our 129 datasets and then averaged these maximum levels across transcripts. The results indicate that lncRNAs are expressed at lower levels than coding RNAs (Figure 2A), with coding regions expressed at three-fold higher levels, on average, than non-TE-lncRNAs (average FPKM: 12.57 vs. 4.30) and six-fold higher levels, on average, than TE-lncRNAs (average FPKM: 12.57 vs. 2.04).

**Figure 2:**
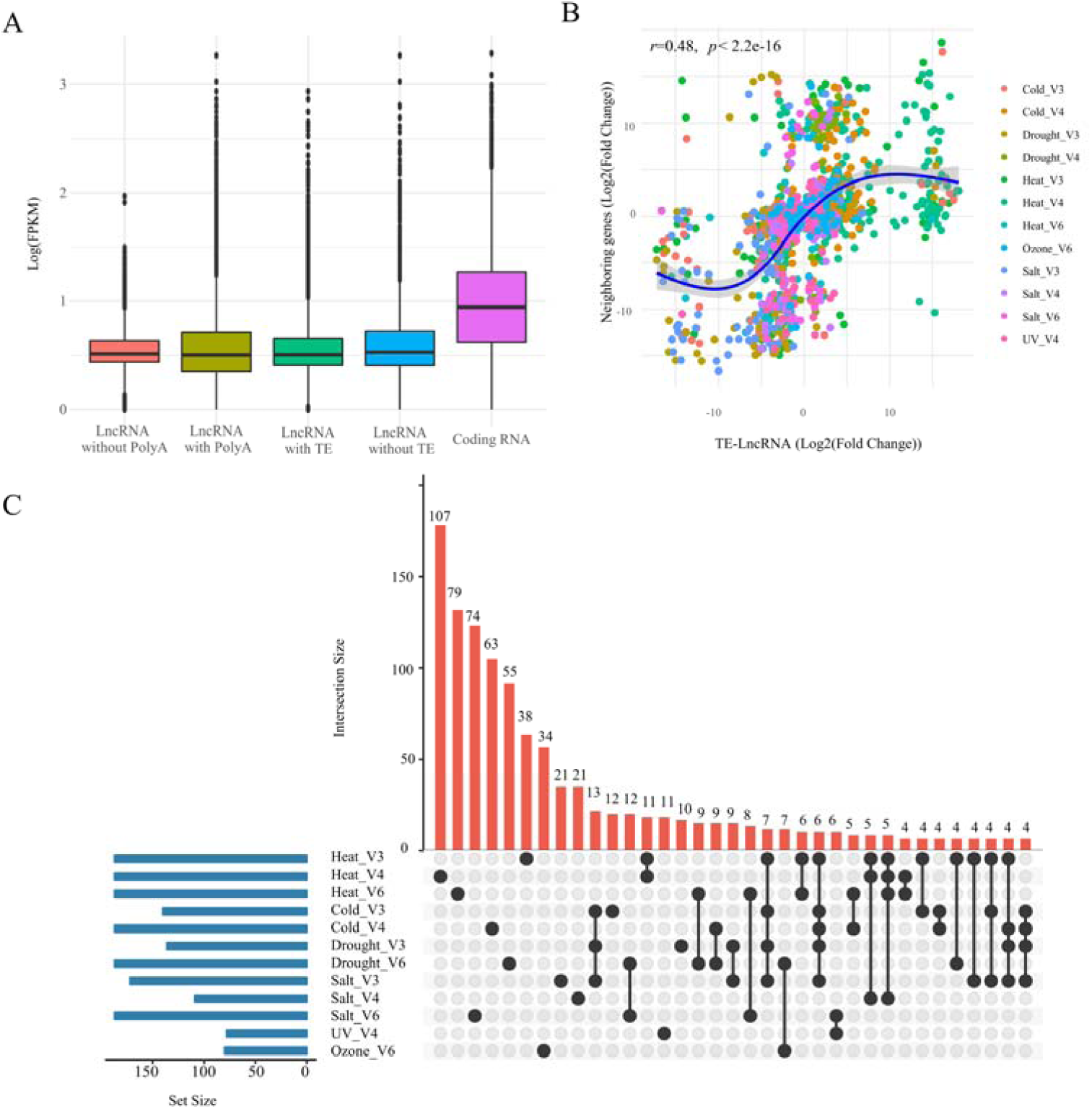
Features of the expression dynamics of lncRNAs. A) The Log2(FPKM) expression level of coding RNAs, lncRNAs and TE-lncRNAs, based on the maximum expression of each lncRNA across datasets. B) The correlation (log2(Fold Change)) of TE -lncRNAs and the closest neighboring gene under different stress conditions. The blue curve indicates the best fit across all of the plotted points and clearly indicates a strong positive correlation from when log2(Fold Change) varies between roughly −5 and 5. The linear correlation of *r*=0.48 is indicated in the graph. C) This figure reports the number of differentially expressed lncRNAs under different stress conditions and developmental stages. The blue histogram on the left shows how many lncRNAs were differentially expressed under different conditions. The red histogram, coupled with the dot plot below, represent the distribution of differentially expressed lncRNAs among stress and stages. For example, the blue graph on the left indicates that 187 lncRNAs were differentially expressed under heat stress in V3. Of these, 38 were detected only under heat stress, as indicated by the graph in red, while 21 were differentially expressed under heat stress at both the V3 and V4 stages and 11 were differentially expressed under heat stress at both the V3 and V6 stages.

We next sought to identify coding genes and lncRNAs that were differentially expressed under abiotic stress. To do so, we contrasted treatment versus control RNAseq samples. For example, the RNAseq data from V3 consisted of two control samples and two replicated treatment samples from each of four stresses (salt, drought, heat and cold) (Table S1). Accordingly, we contrasted each stress treatment to the control, for a total of four contrasts in the V3 stage. Extending this approach to the V4 and V6 stages across all the RNAseq data, we performed a total of 12 contrasts (Table S1). These contrasts identified numerous differentially expressed coding genes and lncRNAs (Table 3). The various treatments identified ∼2000 up- or down-regulated coding transcripts, on average, and a set of 1,077 non-redundant lncRNAs that were either up- and down-regulated across treatments.

**Table 3:** Numbers of differentially expressed genes, lncRNAs and TE-lncRNAs in maize seedlings under abiotic stress.

| Condition/Stage | Heat |  |  | Cold |  | Drought |  | Salt |  |  | UV | Ozone |
| --- | --- | --- | --- | --- | --- | --- | --- | --- | --- | --- | --- | --- |
|  | V3 | V4 | V6 | V3 | V4 | V3 | V6 | V3 | V4 | V6 | V4 | V6 |
| **Up-regulated** | 2,051 | 3,279 | 2,892 | 1,273 | 3,161 | 1,260 | 2,560 | 1,292 | 2,348 | 3,114 | 2,752 | 1,712 |
| Coding | 1,952 | 3,030 | 2,681 | 1,248 | 2,976 | 1,233 | 2,433 | 1,264 | 2,264 | 3,028 | 2,703 | 1,660 |
| lncRNA | 84 | 223 | 181 | 23 | 159 | 24 | 112 | 22 | 64 | 74 | 36 | 45 |
| TE-lncRNA | 36 | 92 | 86 | 6 | 64 | 8 | 59 | 5 | 21 | 31 | 13 | 19 |
| **Down-regulated** | 1,511 | 3,411 | 3,407 | 1,450 | 2,420 | 1,543 | 1,904 | 1,266 | 2,944 | 3,183 | 2,740 | 738 |
| Coding | 1,395 | 3,299 | 3,209 | 1,312 | 2,310 | 1,414 | 1,806 | 1,101 | 2,894 | 2,945 | 2,686 | 699 |
| lncRNA | 103 | 97 | 101 | 117 | 93 | 112 | 80 | 150 | 45 | 208 | 42 | 35 |
| TE-lncRNA | 66 | 38 | 118 | 70 | 39 | 63 | 38 | 101 | 15 | 133 | 14 | 15 |

Among the 1,077 non-redundant lncRNA transcripts, many were differentially expressed in two or more treatments. For example, 679 lncRNAs were identified as differentially expressed across V3-V6 stages under heat treatment (Table 3; Figure 2C; Table S3, Figure S3). Of these, 29 lncRNAs were differentially expressed in all three developmental stages, and 79, 214 and 232 lncRNAs were specific to the V3, V4 and V6 stages, respectively. Interestingly, 40.50% (32/79) heat-responsive lncRNAs at V3 stage, 26.17% (56/214) heat-responsive lncRNAs at V4 and 42.67% (99/232) heat-responsive lncRNAs at V6 were also differentially expressed in response to other stress treatments, but not shared among developmental stages. These patterns implicate many of the lncRNAs in general abiotic stress responses, but they also imply that these responses have temporal (i.e., developmental) specificity.

Interestingly, 529 non-redundant TE-lncRNAs were differentially expressed under one or fewer conditions. The proportion of differentially expressed TE-lncRNAs was lower than the proportion of all lncRNAs; TE-lncRNAs were 65% of the total proportion of lncRNAs, but constituted only 45% and 56% of up- and down-regulated lncRNAs. Most of the differentially expressed TE-lncRNAs had similarity to LTR/*Gypsy* and LTR/*Copia*, as expected, but other TE families also contributed to differentially expressed TE-lncRNAs. For example, MSTRG.32907 exhibited similarities to LINE elements, MSTRG.73329 was similar to DNA/hAT-Ac elements, and MSTRG.37644 was an LTR/Gypsy elements. All of these were differentially expressed in the V3 stage, but in different treatments (heat, cold and salt, respectively) (Figure 4).

lncRNAs have been shown to be involved in *cis* regulation of neighboring genes. To investigate this possibility, we examined the correlation in expression between lncRNAs and their closest neighboring gene in either the 5’ or 3’ direction, yielding a dataset of 1077 differentially expressed lncRNAs and their neighboring genes. The lncRNAs were strongly (*r*=0.48), and highly significantly (*p*< 2e-16) correlated with the expression of their closest neighboring gene (Figure 2B), suggesting that lncRNAs may either be involved in *cis* regulation or are subject to some of the same *cis* regulatory features as their neighboring genes.

### Co-expression modules associated with stress responses

Compared to coding genes and microRNAs, the function of lncRNAs in abiotic stress response remains largely unknown. Computational construction of gene co-expression networks can be a valuable tool for linking lncRNAs and coding RNAs and also for beginning to infer potential biological functions, because co-expressed genes are often members of the same pathway or protein complexes, are often either functionally related, or are controlled by the same transcriptional regulatory program [33, 40–42].

We used the 89 Illumina RNA-seq datasets to build co-expression networks (see Methods and Table S1). WGCNA analyses identified 40 modules that comprise various nodes in the network. Of the 40 inferred modules, 16 were significantly correlated with stress treatments (Figure 3, Figure S4, Table S3, S4). These 16 contained 7,221 transcripts including 408 lncRNAs, of which 171 were TE-lncRNAs. Most of the 16 modules were associated with a single stress and developmental stage, but some were correlated with two or more stresses or stages (Figure 3). For example, the ME_darkgreen module was highly correlated with drought at the V3 stage (*r*^2^=0.76, *p*<4e-18), but it was also significantly correlated with salt stress at the V3 (*r*^2^=0.21, *p*<0.05) and V4 (*r*^2^=0.29, *p*<0.005) stages. Similarly, the ME_salmon module correlated with drought (*r*^2^=0.25, *p*<0.02) and salt stresses (*r*^2^=0.45, *p*<1e-05) and salt stress at the V4 stage (*r*^2^=0.38, *p*<3e-04). Complete correlation information between modules and stress conditions and developmental stages are provided in Figure S4.

**Figure 3:**
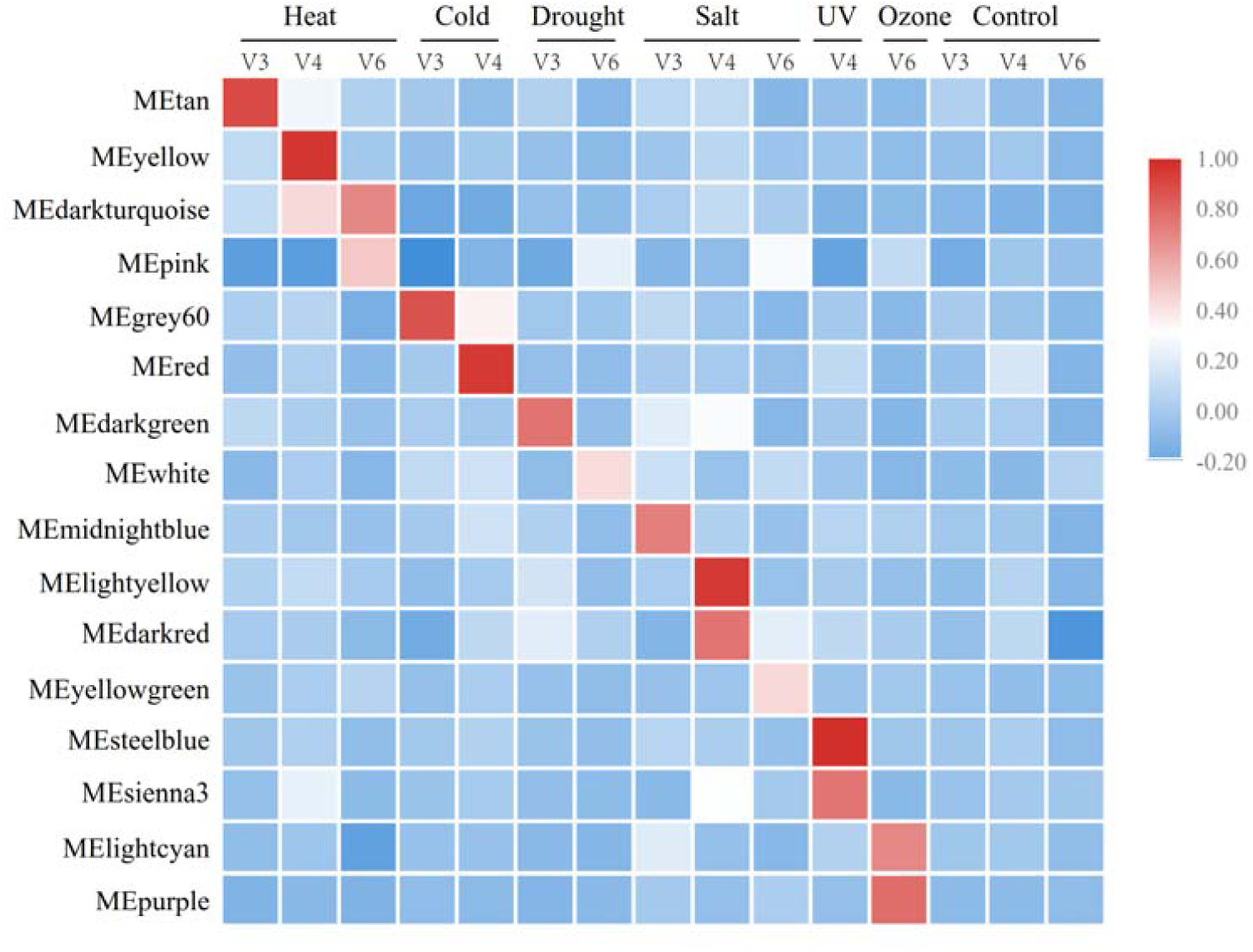
A visual representation of the 16 modules that were significantly correlated with abiotic stress responses. All of the modules were associated with one stress condition and developmental stage, such that they exhibit a temporal cascade of stress responsiveness under different stresses and across V3 to V6 developmental stages. The scale of the heat map reflects the level of correlation (*r*) among genes in an expression module for a specific abiotic stress (i.e., Heat, Cold, Drought, Salt, UV, Ozone) at a specific development stages (i.e., V3 to V6).

**Figure 4:**
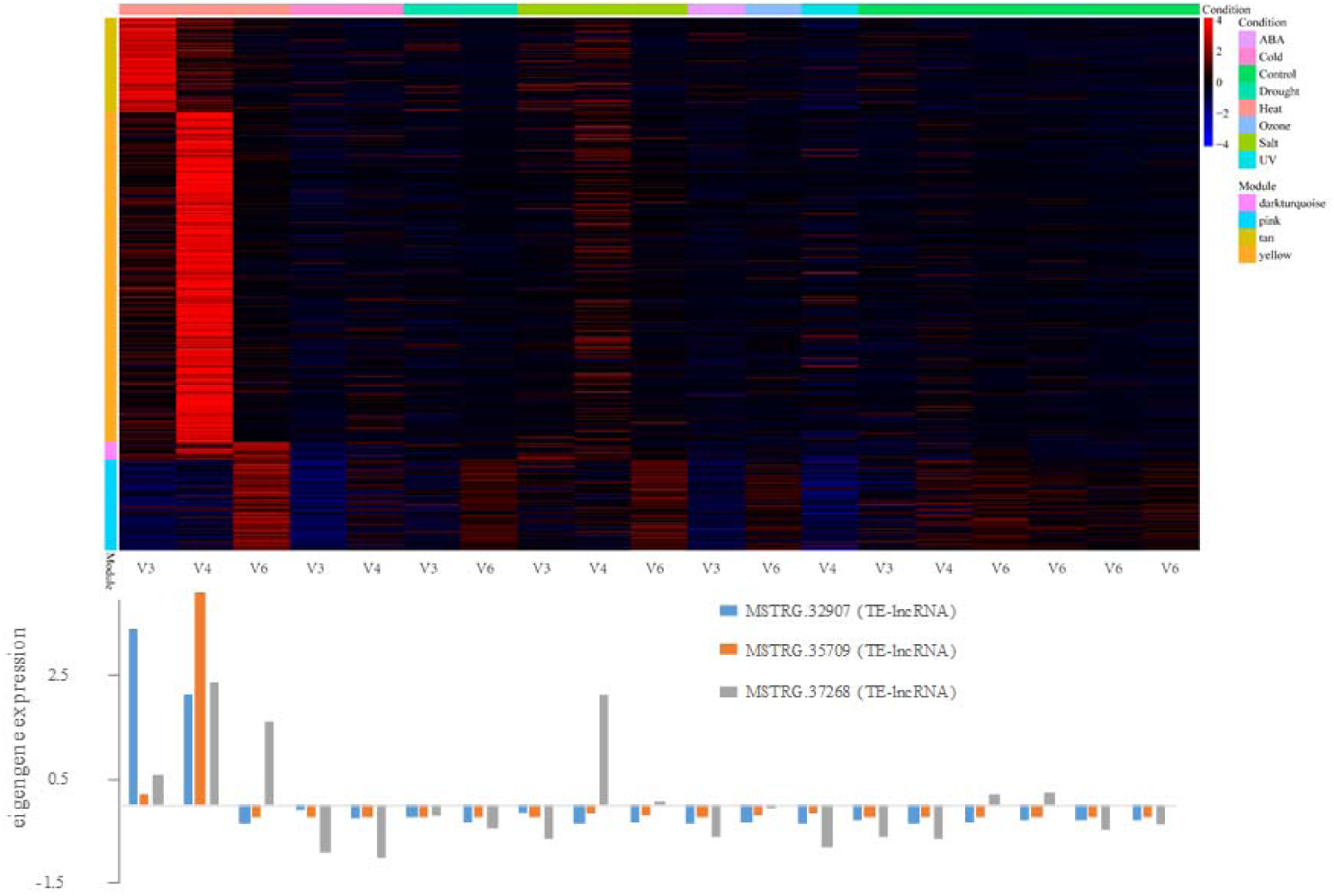
lncRNA expression for four modules associated with heat stress. This figure consists of a heat map (top) and graphs of the expression of specific TE-lncRNAs (bottom) that were chosen because they were top three overrepresented lncRNAs in the four modules and had high interconnectivity. **Top)** The heat graph shows transcript expression levels for hub genes and lncRNAs in each module (*y*-axis) and across conditions (<u>x</u>-axis). The key to modules (*y*-axis) and stress conditions (*x*-axis) are shown on the right legend, with conditions also separated by developmental stage (bottom of *x*-axis). Warmer colors within the heat map indicate high expression, and cooler colors are low (or under) expression. This particular heat map illustrates that the four heat-associated modules are, as expected, highly expressed under heat stress, but not always at the same developmental stage. **Bottom**) The bar plots below the heat graph are eigen-lncRNA expression values selected from the top three overrepresented TE-lncRNAs with high interconnectivity. The x-axis is the same as the heat map, and the id of the TE-lncRNAs is provided by the color key. This graphs shows, again, that the TE-lncRNAs tend to be more highly expressed under heat stress, but with some dependence on developmental stage. Figures S5 to S9 present similar figures for modules associated with cold, drought, salt, UV and ozone stress, respectively.

Recent work uncovered a temporal transcriptional logic underlying nitrogen (N) signaling in *Arabidopsis* [43]; we see similar logic based on developmental timing for abiotic stress responses. Consider the example of heat stress: the ME_tan module was correlated with V3 heat stress (r^2^=0.89, *p*<4e-32), the ME_yellow module correlated with V4 heat stresses (r^2^=0.96, *p*<1e-49), and the ME_darkturquoise (r^2^=0.43, *p*<2e-05) and ME_pink (r^2^=0.49, *p*<1e-06) modules were associated with heat in the V6 stage. These data suggest a developmental cascade of heat-responsive modules. To illustrate this graphically, we arranged the 16 associated modules by stress and development stage. Like heat stress, cold and drought stress were both associated with distinct modules at different developmental stages. There were exceptions, however, as both salt and UV stress associated with two modules in the V4 stage (Figure 3).

Among the 16 significant modules, the most lncRNAs were associated with the ME_yellow module, which correlated with heat stress in the V4 stage (r^2^=0.96, *p*<1e-49) and contained 147 lncRNAs and 65 TE-lncRNAs. Figure 4 details the expression pattern of this and other stress related modules. Given these modules, it is possible to extract the eigengenes from modules to infer function. For example, the eigengenes for the ME_yellow module were assigned into GO categories related to ‘response to heat’, ‘response to high light intensity’, ‘heat acclimation, response to radiation’, ‘regulation of seedling development’ and ‘ER-nucleus signaling pathway’. The ME_darkturquoise (r^2^=0.43, *p*<2e-05) and ME_pink (r^2^=0.49, *p*<1e-06) modules were also associated with heat stress but in a later development stage (V6). These two modules contained 52 lncRNAs and 16 TE-lncRNAs, and their eigengenes exhibited significant enrichment of the GO terms ‘intracellular ribonucleoprotein complex’, ‘HslUV protease complex’, ‘cytoplasmic translation’ and ‘intracellular membrane-bounded organelle’ (Table S5-S7). Overall, GO-inferred functions helped to verify that the modules reflect aspects of the stress response.

### LncRNAs are hubs in modules

An interesting facet of the 16 stress-associated modules is that each contained both lncRNAs and TE-lncRNAs. We have mentioned that the ME_yellow module contained the most lncRNAs of the 16 modules, with 147 lncRNAs and 65 TE-lncRNAs, but other modules were similar in containing lncRNAs. For example, the ME_tan module, which is associated heat stress in V3, contained 26 lncRNAs and 9 TE-lncRNAs. An important question concerns the role of these lncRNAs in expression networks. One role, which is suggested by our results (Figure 2B), is that some of the lncRNAs in modules are co-expressed with genes due to *cis* interactions. It is also possible, however, that lncRNAs regulate genes in *trans*. To investigate this possibility, we screened for key ‘hubs’, which we defined by high connectivity (i.e., intramodular connectivity within the top 10% of all members of the module), membership > 0.9 and high significance (*p*< 0.01) in the module. Based on these filters, we identified 670 hubs that included 39 lncRNAs from different stress-responsive modules (Table S4), of which 18 were TE-lncRNAs.

Considering the heat-responsive modules as an example, the 3 associated modules had 27 lncRNAs as hubs, out of 225 total lncRNAs, with 12 of the 27 categorized as TE-lncRNA. The 27 hub lncRNAs included transcript TE-lncRNAs such as MSTRG.32907 (TE-lncRNA, LINE/L1, *p*< 1.78E-04), MSTRG.35709(TE-lncRNA, LTR/*Gypsy, p*< 2.59E-114), MSTRG.44074 (TE-lncRNA, DNA/hAT-Ac, *p*<2.11E-19) and MSTRG.37268 (TE-lncRNA, DNA/CMC-EnSpm, *p*<1.63E-08). In Figure 4, we illustrate the expression patterns of three of the top-ranked hubs within the heat-stress associated modules, with the top-ranked hubs for the other five abiotic stresses in Figures S5-9. All of these hubs are expressed under stress and demonstrate high intramodular connectivity.

Many hubs in co-expression networks are known to belong to transcription factors (TF) of families such as TCP, AP2/EREBP, MYB, WRKY, NAC, bZIP [44–47]. We found interactions and potential crosstalk between lncRNAs and stress-responsive TFs from these families. In the heat-responsive modules, for example, hub lncRNAs such as MSTRG.32907, MSTRG.36825 and MSTRG.30107 and MSTRG.35709 were connected to TF families such as TCP, NAC, Dof and bHLH, which are known to respond to abiotic stress from previous studies (Figure 5)[48–50].

**Figure 5:**
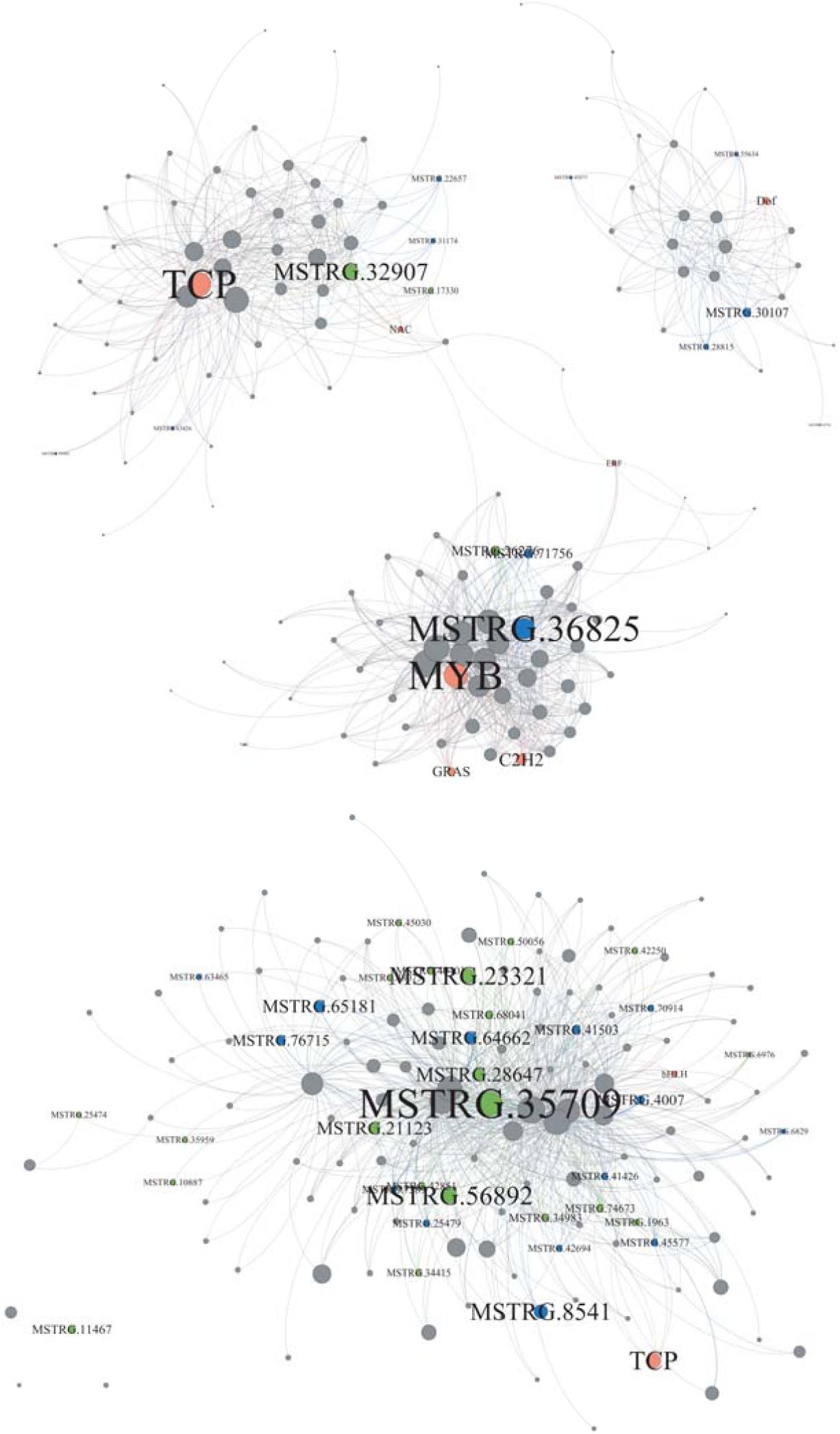
The networks of four heat-responsive modules. The four modules are, the ME_tan module (top left), the ME_yellow module (top right), the ME_pink module (middle), and the ME_darkturquoise module (bottom). In each network diagram, the green circles represent TE-lncRNAs; the blue color represents nonTE-lncRNAs; the orange dots represent known transcription factors from various families (e.g., TCP), and grey circles represent coding RNAs. The size of the dot represents intramodular connectivity, with larger sizes representing higher connectivity. From these networks, we can infer that lncRNAs and TE-lncRNAs are sometimes as or more interconnected than transcription factors.

These results suggest the possibility that lncRNAs – and more specifically, some TE-lncRNAs – act to regulate abiotic stress responses. If they play a functional role, one would expect them to be conserved over evolutionary time. We tested this idea by blasting each of the 39 hub lncRNAs to an evolutionary gradient of genomes that included sorghum, rice and *Arabidopsis* (Table S8). Of the 39, 16 had strong hits (e < 10^−15^) to sorghum, a close relative to maize, and 4 of these 16 were TE-lncRNAs. Three of the hub lncRNAs had hits to rice, but zero of the TE-lncRNAs had rice hits. None of the 39 hub lncRNAs had significant hits to *Arabidopsis*. Overall, these results suggest that ∼10% these lncRNAs have been conserved since the divergence of rice and maize, roughly 50 million years ago [51], and that 39% have been conserved since the divergence between sorghum and maize, roughly 16 million years ago [52].

**Figure 6:**
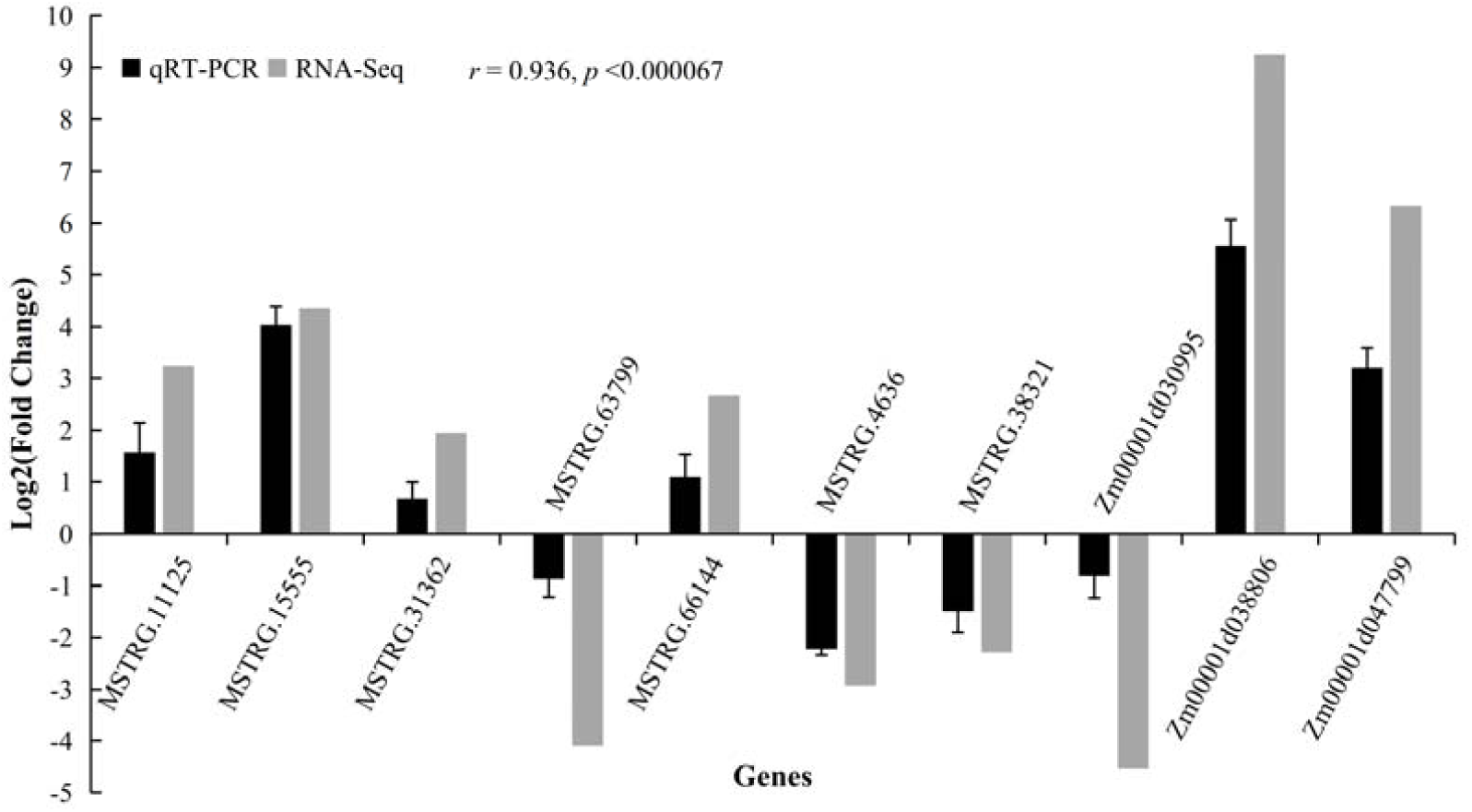
qRT-PCR validation of differential expressed lncRNAs and coding RNAs in RNA-Seq analysis. The qRT-PCR data represents the mean ± standard error (SE) of three independent biological replicates.

### Reliability of gene Expression via qRT-PCR

To verify the reliability of the RNA-seq experiments, a set of independent biological replicates of heat treatment in maize B73 at the stage of three-leaves stage were subjected to quantitative real-time PCR (qRT-PCR) to confirm the expression changes.

Ten transcripts including seven lncRNAs and three coding genes were randomly selected for qPCR. Among them, six genes were significantly up-regulated, four genes were significantly down-regulated under heat stress. The results showed a high degree of consistency for the product sizes between RNA-Seq and qRT-PCR(Figure6, Table S9).

## Discussion

In this study, we accumulated and mined an expansive dataset to identify lncRNAs in maize, particularly those that are expressed in response to abiotic stress. Bioinformatic analyses led to the identification of 23,309 lncRNAs, the largest collection yet identified from maize. We characterized these lncRNAs with respect to three features: *i*) their prevalence and origins, especially lncRNAs that appear to be derived from TEs, *ii*) their expression levels and patterns, including a detectable *cis*-effect, and *iii*) their potential for functioning in abiotic stress response, as inferred from the construction of co-expression networks.

### lncRNA identification and characterization

By its very nature, lncRNA discovery is limited by a number of factors. It is first, of course, limited by the definition of lncRNAs that have been used in the literature – i.e., an RNA molecular > 200 bp with at most one ORF or overlapping exon of < 100 codons [1]. Following precedence, we have adopted this definition for lncRNA discovery, but it bears remembering that some of these could in fact be translated because they contain short ORFs. A second limitation is the fact that our search strategy did not include lncRNAs that overlapped with (or contained) an annotated exon. We applied this limitation purposefully, to avoid mis-classification based on fragmented RNA molecules or contigs. For that reason, however, our work likely underrepresents lncRNAs derived from genes and so some of our estimates may be inaccurate. For example, if many lncRNAs are derived from genic regions, then our estimate of the proportion of lncRNAs that are derived from TE-lncRNAs is an overestimate. It is worth noting, however, that our estimate of the proportion of TE-lncRNAs (65%) is similar to a previous maize study that estimated 68% of lncRNAs were derived from TEs [8]. A third limitation is that the completeness of lncRNA discovery relies critically on the number of tissue and developmental samples that are available. With the exception of *A*. *thaliana*, for which lncRNA discovery was based on 2000 microarray transcriptomes, most plant studies have been limited to only a handful of samples, suggesting that there is still much to learn about the lncRNA complement within and among plant species. To date, the most RNAseq samples use for lncRNA discovery in maize has been 30 [8]; hence, our study has greatly expanded lncRNA discovery in this important crop.

Our RNA datasets were highly enriched for polyadenylated (polyA+) transcripts, because it consisted of 36 PacBio fl-cDNA datasets, 89 RNAseq datasets and only two total RNA datasets. Nonetheless, fully 44% of intergenic lncRNAs were identified from the total RNA data, representing a disproportionately large number relative to polyA+ data. This observation superficially suggests that far more lncRNAs are polyA-, which is an important point to consider when one considers that most – but not all [6, 53, 54] – lncRNA surveys in plants have relied solely on RNAseq samples and not total RNA samples. Previous work has also suggested that the ratio of polyA- and polyA+ lncRNAs may be a function of growth conditions and external stresses (Yuan et al., 2018). A fuller understanding of lncRNAs may require more substantial investments in total RNA datasets.

### Most lncRNAs are TE-lncRNAs

Given our identification of 23,309 lncRNAs, we next sought to characterize their loci of origin and particularly to identify those that likely originated from TEs. We found that ∼65% (15,312) of lncRNAs contained similarity to known TEs. Of these, most (61%, 9341 of 15312) were similar to TEs over >90% of their length, suggesting they were derived solely from TEs. As we noted above, our estimates of the proportion of TE-lncRNAs could be too high, based on our search strategy. However, it is also not surprising that we identified a high proportion of TE-lncRNAs, for at least three reasons. First, previous studies in mammals have demonstrated that most lincRNAs derive from TEs [7, 36]. Second, the maize genome is replete with TEs, with >85% of the genome estimated to consist of DNA derived from TEs [32]. Finally, an important function of lncRNAs is to be precursors for small RNAs, which in turn contribute to TE silencing via sequence homology [8, 55–57].

We also investigated the TE families from which TE-lncRNAs originated. Most of the TE-lncRNAs were derived from LTR/*Gypsy* and LTR/*Copia* families (Table 2), reflecting their preponderance in the maize genome [32, 57]. lncRNAs derived from LTR/*Gypsy* elements were represented in a similar proportion to their genomic proportion (by length) among the TEs we investigated in our study (Table 2). However, LTR/*Copia* elements were underrepresented in the TE-lncRNA dataset relative to their combined lengths in the genome, 22% versus 33%. This suggests that LTR/*Copia* elements do not produce lncRNAs as readily as LTR/*Gypsy* elements, at least within our data. The reasons for the difference between LTR/*Copia* and LTR/*Gypsy* are presently unclear, but one can consider two broad categories: TE age and TE location. For the former, older elements might be expected to be in a deeply-silenced epigenetic state that relies primarily on the maintenance of methylation during cell division rather than an active epigenetic response that enlists lncRNAs[58]. For the latter, one might expect LTR/*Copia* elements to be in genomic locations that are transcribed. In fact, however, the opposite is true, because LTR/*Gypsy* elements tend to be concentrated in pericentromeric regions [32] where there may be less active transcription and less ongoing silencing. In contrast, LTR/*Copia* elements tend to accumulate preferentially in euchromatic regions[32] that tend to be more transcriptionally active. Class II DNA elements also tend to be located near genes and euchromatic regions, but unlike LTR/*Copia* elements they produce lncRNAs at about a 2-fold higher than implied by their genomic lengths (Table 2). To sum: we have shown that TE superfamilies over- and under-produce lncRNAs relative to their genomic representation based on our extensive collection of datasets, but the ultimate causes of these differences remain unclear.

### Levels and patterns of lncRNA expression

Several previous papers from both plants and animals have shown that lncRNAs tend to be expressed at lower levels than *bona fide* genes and that they also tend to show tissue-specific patterns of expression [3, 7–12]. We have verified the former by recording the maximum FPKM for each lncRNA transcript across datasets; on average, lncRNAs are expressed at 4-fold lower levels than genic transcripts by this metric (Figure 2A). Unfortunately, we cannot verify that lncRNAs have more tissue specific expression than genes, because the bulk of our data were isolated from leaves. We can, however, verify that they have lower entropy than genes, on average (Average Shannon Entropy = 2.10 for coding genes vs. 1.13 for lncRNAS), because the lncRNAs consistently lack expression evidence under more conditions.

Of the 13,822 polyA+ lncRNAs, we found that 1,077 (7.79%) were differentially expressed under stress conditions, including 529 TE-lncRNAs. These TE-lncRNAs provided an opportunity to assess whether they could be linked to the expression of nearby genes, indicating some sort of *cis*-regulatory pattern, as has been observed in other species [20, 59, 60]. TE-lncRNAs were significantly correlated (r^2^=0.48; *p*< 2.0e-16) with their nearest neighboring genes (Figure 2B), suggesting that TE-lncRNAs may either be involved in *cis* regulation or are subject to some of the same *cis* regulatory features as their neighboring genes, such as open chromatin structure.

### lncRNAs, abiotic stress and coexpression modules

This study was designed specifically to identify stress-responsive lncRNAs. We approached this problem in two ways. We first identified differentially regulated lncRNAs from a series of controlled experiments for heat, cold, drought and salt stress. Comparing the stress treatment to their corresponding control across 12 different contrasts, we identified 1,077 lncRNAs with evidence for differential expression. This observation corroborates previous studies in suggesting that lncRNAs may be differentially regulated under stress [6, 22–24, 30, 31], but it provides no indication whether the differentially regulated lncRNAs are a byproduct of stress responses or play a functional role. There is, however, a large gap between observing differential expression and proving function. As a first step toward bridging this gap, we have built co-expression networks based on both coding RNAs and lncRNAs from 89 RNAseq datasets, yielding a total of 40 co-expression modules. Of these, 16 were significantly associated with stress responses, and GO annotations of these modules were generally consistent with their inferred response functions. One interesting facet of these 16 modules is that they demonstrate clear patterns across developmental time (Figure 3), suggesting that temporal hierarchies are important for plant responses to environmental stress.

It is difficult to infer function from co-expression modules [61], but studies have shown that genes with high connectedness tend to be functionally essential [62, 63]. We were therefore particularly interested whether any of our lncRNAs are included within co-expression networks and particularly whether they are ‘hubs’ within network modules. Of the 16 modules that were significantly associated with stress responses, we identified 670 hubs, many of which corresponded to genes from known transcription factor families (Figure 5). Of these 670 hubs, 39 were lncRNA transcripts. These represent our best candidates for lncRNAs that function in stress response, potentially as *trans*-acting regulatory factors. Consistent with this last conjecture, several of these lncRNA hubs were connected to genes from known TF factors [48–50]. Moreover, ∼10% these lncRNAs yield strong blast hits to rice, suggesting some measure of evolutionary conservation consistent with functional constraint, at least for this subset.

One somewhat surprising finding is that 18 of the 39 lncRNA hubs are related in sequence to – and perhaps derived from - TEs. This observation raises the intriguing idea that TE exaptation can occur at the level of lncRNAs. It is now well known that TE exaptation contributes to many aspects of genome function, including protein coding genes and especially functional regulatory elements [64–66]. The location of TE-lncRNAs as hubs, along with their connectedness to known TFs, suggests that a small subset of TE-derived lncRNAs may function as *trans*-acting regulatory factors in maize. If true, these hubs appear to have been recruited recently, given that only four of 16 yield strong hits to the sorghum genome. Clearly additional work is required to prove that these TE-lncRNAs function as hypothesized in abiotic response, but their centrality in co-expression modules is nonetheless an intriguing result that is consistent with previous findings showing that most lncRNAs are derived from TEs [7] and that lncRNAs can play central regulatory roles in plant and animal development [64].

## Methods

### Sample collection

In this study, we gathered 36 Pacbio (Pacific Biosciences) Isoseq datasets that were sampled from different tissues [67] and 91 illumina RNAseq datasets that were sampled from leaves of maize B73 [6, 68–70] (Table S1, Figure S1). Of the Illumina datasets, 89 represented polyA+ transcripts and two were based on total RNA, which includes putative polyA-transcripts. The datasets were used for three purposes: lncRNA discovery, differential gene expression analyses, and the inference of gene co-expression networks. All of the 127 datasets were used for lncRNA discovery. A subset of 71 of the 91 RNAseq datasets were employed for differential gene expression analyses (Table S1); these included replicated control and treatment samples from experiments that tested the effects of drought, salt, heat, cold, UV and ozone treatments on gene expression. Finally, all of the 89 polyA+ illumine RNAseq datasets were used for inferring gene co-expression networks. The 89 Illumina datasets represented a developmental series sampled from V3, V4 and V6 seedlings; we take advantage of this developmental series in some network analyses (Table S1, Figure S1).

### Data processing and alignment

Raw data were converted into the FASTQ-formatted file by the Fastq-dump program from the SRA Toolkit (https://github.com/ncbi/sratoolkit). For Illumina data, the SolexaQA++ v3.1 program [71] was employed for quality trimming, using the Q20 value. After trimming, any reads < 50 bp were removed. Cleaned reads were then aligned to the B73 reference genome sequence (v4, http://plants.ensembl.org) using the STAR aligner program [72] with default parameters. Aligned reads were assembled into transcripts by the StringTie program, using the RABT (reference annotation-based transcript) assembly algorithm [73]. For the Pacbio IsoSeq data, reads were aligned to the B73 reference genome using the Minimap2 program [74]. Unique isoforms were collapsed, based on genome alignment by Cupcake ToFU (https://github.com/Magdoll/cDNA_Cupcake). Subsequently, the assembled transcripts from Illumina RNAseq and PacbioIsoSeq were merged using StringTie, which yielded a non-redundant unified set of transcripts.

### Computational identification of intergenic and intronic lncRNAs

To find lncRNAs, a strict computational strategy was performed as described by Lv et. al (2016) that and consisted of four steps. First, non-redundant transcripts were submitted to annotation programs to evaluate their coding potential. We used two annotation programs – CPC2[75]and CNCI[76] – and focused on transcripts that were identified as having no coding potential by both programs as candidate lncRNAs. Second, we submitted candidates to the Pfam database using Pfam_scan script (ftp://ftp.ebi.ac.uk/pub/databases/Pfam/), which aligns transcripts with Hmmer[77]. We filtered out any transcripts that aligned to known protein families at an Evalue<1e-05. Third, we compared the remaining transcripts to reference annotations using gffcompare[78], which outputs various codes to designate the relationship of transcripts to annotated coding regions. We retained transcripts with class codes “i”, which indicates that a transcript is fully contained within a reference intron, and “u”, which designates transcripts that are not obviously related to known coding regions, for further analyses. This last step is likely to miss some sense and anti-sense lncRNAs that derive from coding regions but also limit false positives based on incompletely assembled coding transcripts. Finally, we retained transcripts as high confidence lncRNAs if they passed all of the previous four steps, if they were longer than 200bp, and if they had an FPKM (fragments per kilobase of exon model per million reads mapped) > 1 in at least one of our sample datasets. To determine the relationship of high-confidence lncRNAs to TEs, we masked the lncRNA sequences to identify TE domains. Masking was based on the maize-specific library of Repbase database (www.girinst.org) and was performed by RepeatMasker (www.repeatmasker.org).

### Gene expression analyses

We performed two separate types of analyses based on gene and lncRNA expression data. The first analysis was differential expression analysis based on comparisons between stress and control data (Table S1). To perform these analyses, high quality reads were aligned to the B73 reference using the STAR program [72]. For reads that mapped to multiple locations, we removed alignment reads with a mapping quality <20, based on SAMTools [79]. Raw counts were quantified using the featureCounts program [80], and the FPKM value per gene was calculated using a custom Perl script. The DESeq2 package [81] was used to perform pairwise comparisons between samples to identify differentially expressed transcripts. To identify differentially expressed genes (DEG), we relied on two criteria: the Log2(fold change) had to be >1 and the adjusted p-value from DEseq analyses had to be *p*-adj<0.05.

The second type of analysis was the inference of co-expression networks. To construct networks, expression profiles were extracted from each gene and lncRNA, and expression levels were normalized using variance stabilizing transformation in DESeq2 [81]. Co-expression correlations among lncRNAs and genes were based on Pearson correlations with R^2^⍰≥⍰0.8 across the 89 RNAseq datasets. An unsigned co-expression network was inferred using the WGCNA package[82] with an optimal soft threshold⍰=⍰12. Modules within the network were assigned using Topological Overlap Matrix (TOM). The correlations between modules and stress treatments were calculated and plotted, and then the significant stress-responsive modules were extracted for further analysis.Co-expressed networks were visualized by the Gephi program [83].

### Gene Ontology enrichment analysis

The eigengene probes of each stress-responsive module were assigned putative functions by searching against the UniProt protein database [84]. Searching was based on using the Blastx program [85], using a cut-off e-value⍰≤⍰1e-10. Coding eigengenes were then submitted to the AgriGO v2 online toolkit [86] for gene ontology term enrichment. A Fisher’s exact test was applied for the enrichment analysis and the *p* value was adjusted using the Bonferroni method, with an experiment-wide significance level of 0.05.

### Stress treatment, RNA extraction and qRT-PCR analysis

The maize inbred line B73 was germinated in a greenhouse at JAAS (Jiangsu Academy of Agricultural Sciences). Seedlings at the three-leaf (V3) stage were then incubated at 50°C for 4 hours for heat stress treatment as described by Makarevitch et al. [70]. Leaves with three independent biological replicates were collected and processed for RNA extraction and first strand cDNA synthesis according to PrimeScriptTM^RT^ Master Mix (TaKaRa). qPCR was performed using SYBR Premix DimerEraser™ kits (Takara) on a Real Time PCR System (Roche LightCyclerR 96, USA), according to the manufacturer’s instructions. Quantification results of target transcripts were calculated using the comparative 2ΔΔCt method. Primers were designed using Primer Primer5 [87] and can be found in Additional file 1: Table S9

## Acknowledgements

The authors would like to acknowledge the support of High-Performance Computing Center at UC Irvine.

## Authors’ contributions

BSG and YDL conceived and designed the study. YDL and FQH performed the lncRNA discovery; YFZ and YDL performed the TE annotation. FQH and FLW performed the construction of co-expression network analysis, heat treatment and qRT-PCR experiment. FQH, YDL and BSG wrote the manuscript. All authors read and approved the manuscript.

## Ethics approval and consent to participate

Ethics approval was not required for this study.

## Consent for publication

Not applicable.

## Open Access

This article is distributed under the terms of the Creative Commons Attribution 4.0 International License (http://creativecommons.org/licenses/by/4.0/), which permits unrestricted use, distribution, and reproduction in any medium, provided you give appropriate credit to the original author(s) and the source, provide a link to the Creative Commons license, and indicate if changes were made. The Creative Commons Public Domain Dedication waiver (http://creativecommons.org/publicdomain/zero/1.0/) applies to the data made available in this article, unless otherwise stated.

## Additional files

**Figure S1** A schematic showing the data used in this paper, the bioinformatic pipeline for lncRNA identification, numbers of genes and identified lcRNAs, and some features of downstream lncRNA analyses.

**Figure S2** The chromosomal distribution of lnc RNAs. Density were plotted on each chromosome.

**Figure S3** Differential expressed genes under different abiotic stresses. A-D, Venn diagram of DEGs under different stresses across different development stage V3-V6. E-G, Venn diagram of DEGs under different stress conditions at one stage.

**Figure S4** The correlation values (*r*) between each of the inferred coexpression modules and the specific traits (e.g., Heat, Cold) at different development times (e.g., V3 to V6). A subset of these data for the top 16 stress-associated modules is provided as a heat map in Figure 3.

**Figure S5-S9** These figures are the same format as that of Figure 4, but represent cold (Figure S5), drought (Figure S6), salt (Figure S7), UV (Figure S8) and ozone stress (Figure S9). Each figure contains a heat map (top) and graphs of the expression of specific TE-lncRNAs (bottom) that were chosen because they overrepresented with high interconnectivity. The heat graph shows transcript expression levels for genes and lncRNAs in each module (y-axis) and across conditions (x-axis). The key to modules (y-axis) and stress conditions (x-axis) are shown on the right legend, with conditions also separated by developmental stage (bottom of x-axis). Warmer colors within the heat map indicate high expression, and cooler colors are under-expression. The bar plots below the heat graph are eigen-lncRNA expression values selected from the top overrepresented lncRNAs with high interconnectivity. The x-axis is the same as the heat map, and the id of the lncRNAs is provided by the color key.

**TableS1** The summary of pre-processing, alignment of Illumina and Pacbio datasets. Table S2 The detail of identified lncRNAs including TE information

**Table S3** Differential expressed lncRNAs and TE-lncRNAs under different abiotic stresses across developmental stages. Each sheet represents different stress treatment at the developmental stage.

**Table S4** The information of stress-associated modules.

**Table S5** The detail of stress-responsive modules including member id, TE superfamily and TF family classification, membership value, kIM value and p value across different stages and stress treatments.

**Table S6** Overrepresented GO terms of stress-responsive modules.

**Table S7** GO enrichment results of different stress-responsive modules. Each sheet include enriched GO term under difference abiotic stress such as heat, cold, drought, salt, UV and Ozone across V3 to V6 stages.

**Table S8** The similarity of 39 hub lncRNAs with neighboring species

## References

1. Mercer TR, Dinger ME, Mattick JS. Long non-coding RNAs: insights into functions. Nat Rev Genet. 2009;10:155–9.

2. Maeda N, Kasukawa T, Oyama R, Gough J, Frith M, Engström PG, et al. Transcript annotation in FANTOM3: mouse gene catalog based on physical cDNAs. PLoS Genet. 2006;2:e62.

3. Derrien T, Johnson R, Bussotti G, Tanzer A, Djebali S, Tilgner H, et al. The GENCODE v7 catalog of human long noncoding RNAs: analysis of their gene structure, evolution, and expression. Genome Res. 2012;22:1775–89.

4. Liu J, Jung C, Xu J, Wang H, Deng S, Bernad L, et al. Genome-Wide Analysis Uncovers Regulation of Long Intergenic Noncoding RNAs in Arabidopsis. Plant Cell. 2012;24:4333–45.

5. Zhang Y-C, Liao J-Y, Li Z-Y, Yu Y, Zhang J-P, Li Q-F, et al. Genome-wide screening and functional analysis identify a large number of long noncoding RNAs involved in the sexual reproduction of rice. Genome Biol. 2014;15:512.

6. Lv Y, Liang Z, Ge M, Qi W, Zhang T, Lin F, et al. Genome-wide identification and functional prediction of nitrogen-responsive intergenic and intronic long non-coding RNAs in maize (Zea mays L.). BMC Genomics. 2016;17:350.

7. Kapusta A, Kronenberg Z, Lynch VJ, Zhuo X, Ramsay L, Bourque G, et al. Transposable elements are major contributors to the origin, diversification, and regulation of vertebrate long noncoding RNAs. PLoS Genet. 2013;9:e1003470.

8. Li L, Eichten SR, Shimizu R, Petsch K, Yeh C-T, Wu W, et al. Genome-wide discovery and characterization of maize long non-coding RNAs. Genome Biol. 2014;15:R40.

9. Zhang H, Hu W, Hao J, Lv S, Wang C, Tong W, et al. Genome-wide identification and functional prediction of novel and fungi-responsive lincRNAs in Triticum aestivum. BMC Genomics. 2016;17:238.

10. Wang M, Yuan D, Tu L, Gao W, He Y, Hu H, et al. Long noncoding RNAs and their proposed functions in fibre development of cotton (Gossypium spp.). 2015;:1181–97.

11. Wang H, Chung PJ, Liu J, Jang IC, Kean MJ, Xu J, et al. Genome-wide identification of long noncoding natural antisense transcripts and their responses to light in Arabidopsis. Genome Res. 2014;24:444–53.

12. Golicz AA, Singh MB, Bhalla PL. The long intergenic noncoding RNA (lincRNA) landscape of the soybean genome. Plant Physiol. 2018;176:2133–47.

13. Yuan J, Li J, Yang Y, Tan C, Zhu Y, Hu L, et al. Stress-responsive regulation of long non-coding RNA polyadenylation in Oryza sativa. Plant J. 2018;93:814–27.

14. Cho J. Transposon-Derived Non-coding RNAs and Their Function in Plants. Front Plant Sci. 2018;9:600.

15. Hadjiargyrou M, Delihas N. The intertwining of transposable elements and non-coding RNAs. Int J Mol Sci. 2013;14:13307–28.

16. Rinn JL, Chang HY. Genome regulation by long noncoding RNAs. Annu Rev Biochem. 2012;81:145–66.

17. Wang T, Zhao M, Zhang X, Liu M, Yang C, Chen Y, et al. Novel phosphate deficiency-responsive long non-coding RNAs in the legume model plant Medicago truncatula. J Exp Bot. 2017;68:5937–48.

18. Kim D-H, Sung S. Vernalization-triggered intragenic chromatin loop formation by long noncoding RNAs. Dev Cell. 2017;40:302-312.e4.

19. Seo JS, Sun H-X, Park BS, Huang C-H, Yeh S-D, Jung C, et al. ELF18-induced long-noncoding RNA associates with mediator to enhance expression of innate immune response genes in Arabidopsis. Plant Cell. 2017;29:1024–38.

20. Kopp F, Mendell JT. Functional classification and experimental dissection of long noncoding RNAs. Cell. 2018;172:393–407.

21. Kung JTY, Colognori D, Lee JT. Long noncoding RNAs: past, present, and future. Genetics. 2013;193:651–69.

22. Ben Amor B, Wirth S, Merchan F, Laporte P, d’Aubenton-Carafa Y, Hirsch J, et al. Novel long non-protein coding RNAs involved in Arabidopsis differentiation and stress responses. Genome Res. 2009;19:57–69.

23. Di C, Yuan J, Wu Y, Li J, Lin H, Hu L, et al. Characterization of stress-responsive lncRNAs in Arabidopsis thaliana by integrating expression, epigenetic and structural features. Plant J. 2014;80:848–61. doi:10.1111/tpj.12679.

24. Liu J, Jung C, Xu J, Wang H, Deng S, Bernad L, et al. Genome-wide analysis uncovers regulation of long intergenic noncoding RNAs in Arabidopsis. Plant Cell. 2012;24:4333–45.

25. Zhu J-K. Abiotic stress signaling and responses in plants. Cell. 2016;167:313–24.

26. Mittler R. Abiotic stress, the field environment and stress combination. Trends Plant Sci. 2006;11:15–9.

27. Masuka B, Araus JL, Das B, Sonder K, Cairns JE. Phenotyping for abiotic stress tolerance in maize. J Integr Plant Biol. 2012;54:238–49.

28. Gong F, Yang L, Tai F, Hu X, Wang W. Omics of maize stress response for sustainable food production: opportunities and challenges. Omics. 2014;18:714–32.

29. Halford NG, Curtis TY, Chen Z, Huang J. Effects of abiotic stress and crop management on cereal grain composition: implications for food quality and safety. J Exp Bot. 2015;66:1145–56.

30. Zhang W, Han Z, Guo Q, Liu Y, Zheng Y, Wu F, et al. Identification of maize long non-coding RNAs responsive to drought stress. PLoS One. 2014;9:e98958.

31. Singh U, Khemka N, Rajkumar MS, Garg R, Jain M. PLncPRO for prediction of long non-coding RNAs (lncRNAs) in plants and its application for discovery of abiotic stress-responsive lncRNAs in rice and chickpea. Nucleic Acids Res. 2017;45:e183.

32. Schnable PS, Ware D, Fulton RS, Stein JC, Wei F, Pasternak S, et al. The B73 maize genome: complexity, diversity, and dynamics. Science. 2009;326:1112–5.

33. van Dam S, Võsa U, van der Graaf A, Franke L, de Magalhães JP. Gene co-expression analysis for functional classification and gene–disease predictions. Brief Bioinform. 2017;19:bbw139.

34. Signal B, Gloss BS, Dinger ME. Computational approaches for functional prediction and characterisation of long noncoding RNAs. Trends Genet. 2016;32:620–37.

35. Jiao Y, Peluso P, Shi J, Liang T, Stitzer MC, Wang B, et al. Improved maize reference genome with single-molecule technologies. Nature. 2017;546:524.

36. Kelley D, Rinn J. Transposable elements reveal a stem cell-specific class of long noncoding RNAs. Genome Biol. 2012;13:R107.

37. Johnson R, Guigó R. The RIDL hypothesis: transposable elements as functional domains of long noncoding RNAs. RNA. 2014;20:959–76.

38. Tsoi LC, Iyer MK, Stuart PE, Swindell WR, Gudjonsson JE, Tejasvi T, et al. Analysis of long non-coding RNAs highlights tissue-specific expression patterns and epigenetic profiles in normal and psoriatic skin. Genome Biol. 2015;16:24.

39. Wang G, Wang G, Zhong M, Wang J, Zhang J, Tang Y, et al. Genome-wide identification, splicing, and expression analysis of the myosin gene family in maize (Zea mays). J Exp Bot. 2014;65:923–38.

40. Amar D, Safer H, Shamir R. Dissection of regulatory networks that are altered in disease via differential co-expression. PLoS Comput Biol. 2013;9:e1002955.

41. Serin EAR, Nijveen H, Hilhorst HWM, Ligterink W. Learning from co-expression networks: possibilities and challenges. Front Plant Sci. 2016;7:444.

42. Borrill P, Harrington SA, Simmonds J, Uauy C. Identification of transcription factors regulating senescence in wheat through gene regulatory network modelling. Plant Physiol. 2019;:pp.00380.2019.

43. Varala K, Marshall-Colón A, Cirrone J, Brooks MD, Pasquino A V, Léran S, et al. Temporal transcriptional logic of dynamic regulatory networks underlying nitrogen signaling and use in plants. Proc Natl Acad Sci U S A. 2018;115:6494–9.

44. Mukhopadhyay P, Tyagi AK. OsTCP19 influences developmental and abiotic stress signaling by modulating ABI4-mediated pathways. Sci Rep. 2015;5:9998.

45. Chen L, Han J, Deng X, Tan S, Li L, Li L, et al. Expansion and stress responses of AP2/EREBP superfamily in Brachypodium Distachyon. Sci Rep. 2016;6:21623.

46. Vinocur B, Altman A. Recent advances in engineering plant tolerance to abiotic stress: achievements and limitations. Curr Opin Biotechnol. 2005;16:123–32.

47. Umezawa T, Fujita M, Fujita Y, Yamaguchi-Shinozaki K, Shinozaki K. Engineering drought tolerance in plants: discovering and tailoring genes to unlock the future. Curr Opin Biotechnol. 2006;17:113–22.

48. Singh KB, Foley RC, Oñate-Sánchez L. Transcription factors in plant defense and stress responses. Curr Opin Plant Biol. 2002;5:430–6.

49. Agarwal PK, Jha B. Transcription factors in plants and ABA dependent and independent abiotic stress signalling. Biol Plant. 2010;54:201–12.

50. Lawlor D. Abiotic stress adaptation in plants. Physiological, molecular and genomic foundation. Ann Bot. 2011;107:vii.

51. Gaut BS. Evolutionary dynamics of grass genomes. New Phytol. 2002;154:15–28.

52. Gaut BS, Le Thierry d’Ennequin M, Peek AS, Sawkins MC. Maize as a model for the evolution of plant nuclear genomes. Proc Natl Acad Sci U S A. 2000;97:7008–15.

53. Yang L, Duff MO, Graveley BR, Carmichael GG, Chen L-L. Genomewide characterization of non-polyadenylated RNAs. Genome Biol. 2011;12:R16.

54. Gaiti F, Fernandez-Valverde SL, Nakanishi N, Calcino AD, Yanai I, Tanurdzic M, et al. Dynamic and Widespread lncRNA Expression in a Sponge and the Origin of Animal Complexity. Mol Biol Evol. 2015;32:2367–82.

55. Boerner S, McGinnis KM. Computational identification and functional predictions of long noncoding RNA in Zea mays. PLoS One. 2012;7:e43047.

56. Zhu Q-H, Wang M-B. Molecular functions of long non-coding rnas in plants. Genes (Basel). 2012;3:176–90.

57. Baucom RS, Estill JC, Chaparro C, Upshaw N, Jogi A, Deragon J-M, et al. Exceptional diversity, non-random distribution, and rapid evolution of retroelements in the B73 maize genome. PLoS Genet. 2009;5:e1000732.

58. Bousios A, Gaut BS. Mechanistic and evolutionary questions about epigenetic conflicts between transposable elements and their plant hosts. Curr Opin Plant Biol. 2016;30:123–33.

59. Yan P, Luo S, Lu JY, Shen X. Cis- and trans-acting lncRNAs in pluripotency and reprogramming. Curr Opin Genet Dev. 2017;46:170–8.

60. Kornienko AE, Guenzl PM, Barlow DP, Pauler FM. Gene regulation by the act of long non-coding RNA transcription. BMC Biol. 2013;11:59.

61. Vandepoele K, Quimbaya M, Casneuf T, De Veylder L, Van de Peer Y. Unraveling Transcriptional control in Arabidopsis using cis-regulatory elements and coexpression networks. Plant Physiol. 2009;150:535–46.

62. Jeong H, Mason SP, Barabási A-L, Oltvai ZN. Lethality and centrality in protein networks. Nature. 2001;411:41–2.

63. Barabási A-L, Oltvai ZN. Network biology: understanding the cell’s functional organization. Nat Rev Genet. 2004;5:101–13.

64. Chuong EB, Elde NC, Feschotte C. Regulatory activities of transposable elements: from conflicts to benefits. Nat Rev Genet. 2017;18:71–86.

65. Trizzino M, Park Y, Holsbach-Beltrame M, Aracena K, Mika K, Caliskan M, et al. Transposable elements are the primary source of novelty in primate gene regulation. Genome Res. 2017;27:1623–33.

66. Chuong EB, Elde NC, Feschotte C. Regulatory evolution of innate immunity through co-option of endogenous retroviruses. Science. 2016;351:1083–7.

67. Wang B, Tseng E, Regulski M, Clark TA, Hon T, Jiao Y, et al. Unveiling the complexity of the maize transcriptome by single-molecule long-read sequencing. Nat Commun. 2016;7:11708.

68. Li P, Cao W, Fang H, Xu S, Yin S, Zhang Y, et al. Transcriptomic profiling of the maize (Zea mays L.) leaf response to abiotic stresses at the seedling stage. Front Plant Sci. 2017;8:290.

69. Mimura M, Zallot R, Niehaus TD, Hasnain G, Gidda SK, Nguyen TND, et al. Arabidopsis TH2 encodes the orphan enzyme thiamin monophosphate phosphatase. Plant Cell. 2016;28:2683–96.

70. Makarevitch I, Waters AJ, West PT, Stitzer M, Hirsch CN, Ross-Ibarra J, et al. Transposable elements contribute to activation of maize genes in response to abiotic stress. PLoS Genet. 2015;11:e1004915.

71. Cox MP, Peterson DA, Biggs PJ. SolexaQA: At-a-glance quality assessment of Illumina second-generation sequencing data. BMC Bioinformatics. 2010;11:485.

72. Dobin A, Davis CA, Schlesinger F, Drenkow J, Zaleski C, Jha S, et al. STAR: ultrafast universal RNA-seq aligner. Bioinformatics. 2013;29:15–21.

73. Pertea M, Pertea GM, Antonescu CM, Chang T-C, Mendell JT, Salzberg SL. StringTie enables improved reconstruction of a transcriptome from RNA-seq reads. Nat Biotechnol. 2015; 33(3):290–295.

74. Li H. Minimap2: pairwise alignment for nucleotide sequences. Bioinformatics. 2018;34:3094–100.

75. Wang D, Qu Z, Yang L, Zhang Q, Liu Z-H, Do T, et al. Transposable elements (TEs) contribute to stress-related long intergenic noncoding RNAs in plants. Plant J. 2017;90:133–46.

76. Sun L, Luo H, Bu D, Zhao G, Yu K, Zhang C, et al. Utilizing sequence intrinsic composition to classify protein-coding and long non-coding transcripts. Nucleic Acids Res. 2013;41:e166–e166.

77. Johnson LS, Eddy SR, Portugaly E. Hidden Markov model speed heuristic and iterative HMM search procedure. BMC Bioinformatics. 2010;11:431.

78. Pertea M, Kim D, Pertea GM, Leek JT, Salzberg SL. Transcript-level expression analysis of RNA-seq experiments with HISAT, StringTie and Ballgown. Nat Protoc. 2016;11:1650–67.

79. Li H, Handsaker B, Wysoker A, Fennell T, Ruan J, Homer N, et al. The Sequence Alignment/Map format and SAMtools. Bioinformatics. 2009;25:2078–9.

80. Liao Y, Smyth GK, Shi W. featureCounts: an efficient general purpose program for assigning sequence reads to genomic features. Bioinformatics. 2014;30:923–30.

81. Love MI, Huber W, Anders S. Moderated estimation of fold change and dispersion for RNA-seq data with DESeq2. Genome Biol. 2014;15:550.

82. Langfelder P, Horvath S. WGCNA: an R package for weighted correlation network analysis. BMC Bioinformatics. 2008;9:559.

83. Bastian M, Heymann S, Jacomy M, others. Gephi: an open source software for exploring and manipulating networks. Icwsm. 2009;8:361–2.

84. Apweiler R. UniProt: the Universal Protein knowledgebase. Nucleic Acids Res. 2004;32:115D–119.

85. Altschul S, Madden TL, Schäffer AA, Zhang J, Zhang Z, Miller W, et al. Gapped BLAST and PSI-BLAST: a new generation of protein database search programs. Nucleic Acids Res. 1997;25:3389–402.

86. Tian T, Liu Y, Yan H, You Q, Yi X, Du Z, et al. agriGO v2.0: a GO analysis toolkit for the agricultural community, 2017 update. Nucleic Acids Res. 2017;45:W122–9.

87. Lalitha S. Primer Premier 5. Biotech Software & Internet Report, 2000;1(6):270–272.

